# Nuclear architecture protein Distal antenna balances genome-binding and phase-separation properties to regulate neuroblast competence

**DOI:** 10.1101/2022.06.30.498164

**Authors:** Gillie Benchorin, Maggie Jiaqi Li, Richard Jangwon Cho, Yuxin Hu, Minoree Kohwi

## Abstract

Neural progenitors transit through multiple competence states that restrict production of each neural cell type. In *Drosophila* neuroblasts, a timed genome reorganization relocates the cell fate gene, *hunchback*, to the nuclear periphery, terminating competence to produce early-born neurons. Distal antenna (Dan), a pipsqueak (Psq) superfamily protein, is transiently downregulated at mid-embryogenesis, which is required for this relocation. Here we find that Dan is a highly intrinsically disordered protein, and when its Psq DNA-binding domain is increasingly disrupted, Dan coalesces into steadily larger, interconnected hubs of rapid protein exchange. Consistent with these phase-separation properties, Dan has a predicted LARKS domain, a structural motif that forms reversible interactions associated with phase-separation. In the embryo, loss of either the Psq motif or the LARKS domain abrogates Dan’s ability to maintain neuroblast early competence upon misexpression, suggesting that Dan requires both DNA-binding and phase-separation to regulate neuroblast competence. Finally, we found that Dan strongly interacts with proteins of the nuclear pore complex (NPC), and Elys, a core NPC scaffold protein known to regulate genome architecture, binds the *hb* intron and is required for competence termination. Together, the results support a model for how Dan’s phase-separation properties can mediate dynamic restructuring by balancing genome-binding, self-association, and interaction among nuclear architecture regulators.

## Introduction

The metazoan genome is non-randomly organized, and this organization is thought to underlie cell type specific gene expression programs (Abbott et al., 2005; Lucas et al., 2021; Misteli, 2020; Schoenfelder and Fraser, 2019; Schwartz and Cavalli, 2017). How genome organization contributes to stage-specific cell fate choices by the progenitor is not well known, let alone how it is regulated or restructured over time. These are particularly important questions in brain development, in which a limited pool of progenitors transit through multiple states of competence to produce different neural cell types in a stage-specific manner (Belliveau et al., 2000; Cleary and Doe, 2006; Hirabayashi et al., 2009; Kohwi et al., 2013; Pearson and Doe, 2003). Like their mammalian counterparts, *Drosophila* neuroblasts have limited windows of competence which have the potential to specify particular cell types during development, and their genetic accessibility make them well-suited to study the role of nuclear architecture and trans-acting regulators in neurogenesis (Hafer et al., 2022; Kohwi et al., 2013; Lucas et al., 2021). In the embryonic ventral nerve cord, a bilaterally symmetric set of 30 neuroblasts undergo multiple rounds of self-renewing asymmetric cell divisions. As they divide, they sequentially express a series of temporal identity transcription factors that specify the identities of the neural daughters produced from each division (Doe, 2017; Isshiki et al., 2001; Kohwi and Doe, 2013). Hunchback (Hb), the first of the series, specifies early-born neural fate, and consequently, all early-born neurons actively maintain *hb* transcription as a molecular signature of their birth timing (Grosskortenhaus et al., 2005; Isshiki et al., 2001; Lucas et al., 2021) (Fig. 1A). Neuroblasts typically express Hb for one to two divisions, but maintain competence to generate early-born neurons for several divisions/hours after *hb* is repressed in the neuroblast (Cleary and Doe, 2006; Hafer et al., 2022; Kohwi et al., 2013; Lucas et al., 2021; Pearson and Doe, 2003). Thus, competence reflects the progenitor’s internal state, or its potential, to specify descendent neurons. Competence can be experimentally observed by transgenically misexpressing Hb in the neuroblast and assessing whether the descendant neuron subsequently activates *hb* transcription endogenously, a key molecular signature of early-born identity. Notably, Hb misexpression in the neuroblast does not induce *hb* within the neuroblast, nor does misexpression in the postmitotic neurons induce *hb* expression within the neuron (Grosskortenhaus et al., 2005; Kohwi et al., 2013; Lucas et al., 2021). The neuron can induce *hb* expression only if Hb is present in the neuroblast during the early competence window, and once this period ends, the neuroblast becomes refractory to Hb and can no longer specify early-born identity. Previously, we showed that early competence is terminated when the *hb* gene relocates to the neuroblast nuclear periphery, observed as an increase in the fraction of *hb* loci localized at the nuclear lamina (Hafer et al., 2022; Kohwi et al., 2013; Lucas et al., 2021). Thus, the developmentally-timed change in radial position of the *hb* gene in the neuroblast progenitor “primes” the transcriptional program of the descendent neurons, and its peripheral relocation results in its heritably silenced state. Accordingly, neuroblast competence to specify *hb*-transcribing, early-born molecular identity, provides a biological readout for the neuroblast’s nuclear architecture at the time of the neuron’s specification.

**Figure 1.**
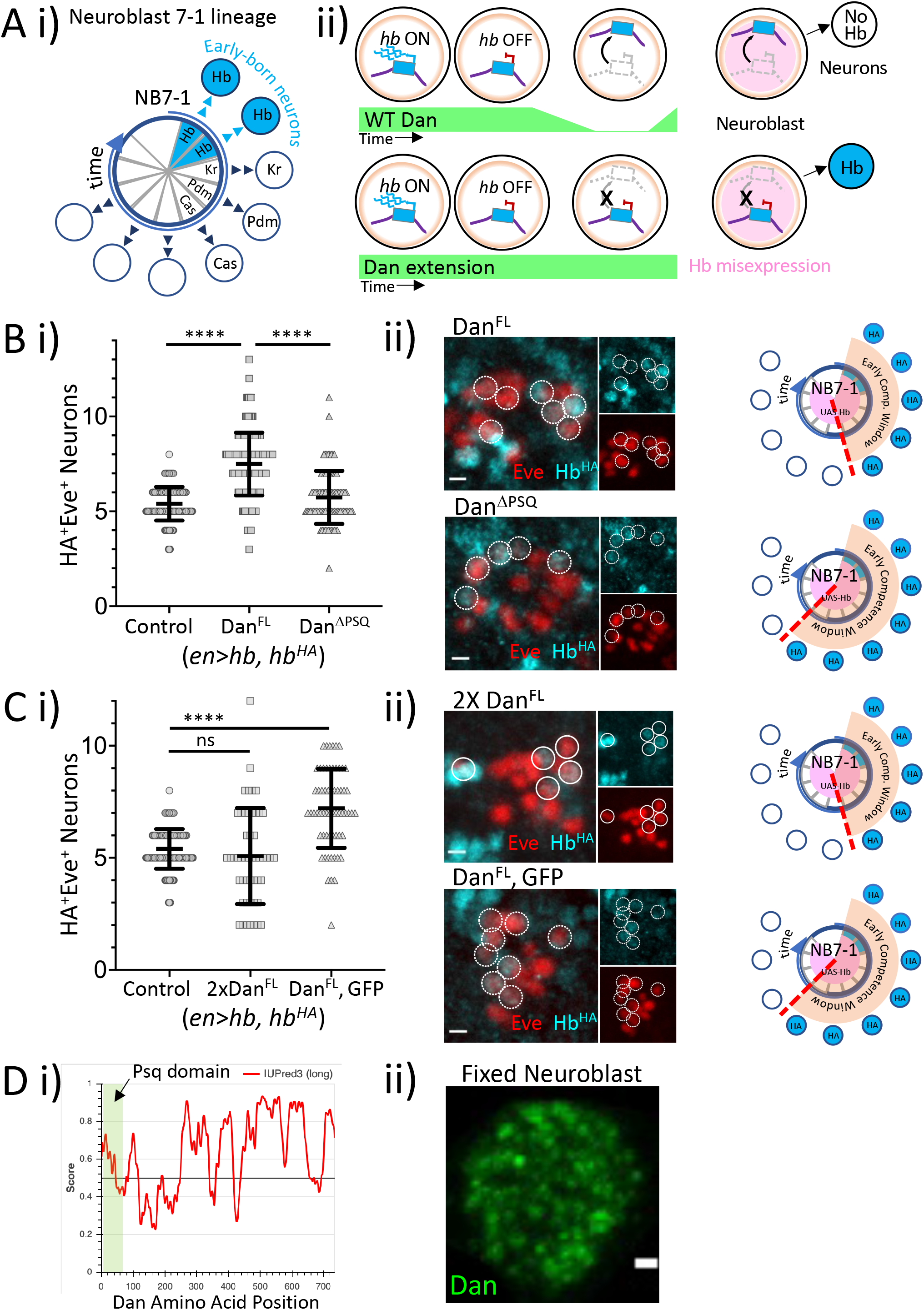
Dan is a highly intrinsically disordered protein and regulates neuroblast competence in a dose-dependent manner. **A) i**, Neuroblast 7-1 (NB7-1) and its neural lineage arranged by birth order. The early-born neurons express Hunchback (Hb). **ii**, Several divisions and hours after *hb* is transcriptionally repressed, it relocates to the nuclear lamina for heritable silencing. This relocation at requires the mid-embryogenic downregulation of Distal Antenna (Dan). Dan misexpression in neuroblasts blocks *hb* relocation and results in extension of early competence. **B) i**, Quantification of the early competence window (Eve^+^HA^+^) comparing control (prolonged Hb only), co-misexpression of Dan^FL^, or Dan^βPSQ^. Each data point represents a single hemisegment, and mean, standard deviation, and t-test significance is shown. **ii**, Images are of single representative hemisegments of the data shown in **i. C)** similar to B, but comparing control to misexpressing two copies of UAS-Dan or one copy each of UAS-Dan and UAS-GFP. Scale bars, 3μm. **D) i**, Dan intrinsic disorder prediction (IUPred3). The Psq domain is highlighted at the N-terminal end. **ii**, Projection of 5 z-planes of a neuroblast immunostained for Dan in whole, fixed stage 9 embryo. Scale bar, 1μm. Also see Supplemental Figure 1.

We previously identified a CNS-specific nuclear factor, Distal antenna (Dan), that is robustly expressed by all neuroblasts at the onset of neurogenesis but is rapidly and transiently downregulated at mid-embryogenesis (Kohwi et al., 2011). This transient downregulation is required for *hb* gene relocation to the neuroblast nuclear lamina, as maintaining Dan expression in neuroblasts through misexpression prevents *hb* relocation and extends the early competence window (Kohwi et al., 2013) (Fig. 1A). Here we show that Dan is a highly disordered protein, and deletion of the Psq motif DNA-binding domain results in Dan protein coalescing into large, liquid droplets. The size of the hubs increases as impairment to the DNA-binding domain increases, suggesting that DNA-binding and self-association create opposing forces on Dan protein distribution within nuclei. Consistently, in addition to the Psq motif DNA-binding domain, Dan has a predicted LARKS domain, a structural motif that forms kinked beta-sheets associated with labile interactions that underlie phase-separation (Hughes et al., 2021). *in vivo*, misexpression of Dan without either the Psq motif or the LARKS domain abrogates Dan’s ability to maintain early competence. Finally, we find that Dan strongly interacts with proteins of the NPC. Among these, we find that Elys, which has been shown to bind DNA and regulate nuclear architecture (Shevelyov, 2020), is required for termination of early competence. The results support a model for how a phase-separating, DNA-binding factor can confer dynamic restructuring to regulate neural progenitor competence transitions.

## Results

### Dan is a highly intrinsically disordered protein and regulates neuroblast competence in a dose-dependent manner

Dan is a member of a large superfamily of proteins whose defining characteristic is the Psq motif helix-turn-helix DNA-binding domain that is highly evolutionarily conserved, spanning insects, vertebrates, nematodes, fungi, and other species (Siegmund and Lehmann, 2002). Within the superfamily, Dan is classified as a member of the CENP-B/Transposase subgroup, which harbors a single Psq domain at the N-terminus and has previously been implicated in the development of the eye and antenna (Curtiss et al., 2007; Emerald et al., 2003). To examine how Dan functions within the neuroblast, we generated a series of UAS constructs to misexpress either full length Dan (Dan^FL^), Dan with a deleted Psq DNA-binding domain (Dan^ΔPSQ^), or the Psq domain fused to GFP (PSQ:GFP), and examined their effects on neuroblast early competence. All transgenes were inserted into identical genomic sites using phi-C31 integration (Venken et al., 2006) (Supp. Fig. 1A) to eliminate differences due to position effects. To perform the competence assay, we used the Gal4/UAS system (Brand and Perrimon, 1993) to drive continuous expression of Hb in the neuroblast using the *engrailed-Gal4* driver, which drives strong expression in row 6 and 7 neuroblasts (Cleary and Doe, 2006; Hafer et al., 2022; Kohwi et al., 2013; Lucas et al., 2021; Pearson and Doe, 2003), and assessed the molecular identity of descendent neurons of NB7-1, a lineage for which we can easily identity the daughter neurons with the expression of the Even-skipped transcription factor (Eve). Given that all early-born neurons are molecularly characterized by active transcription of *hb*, we took advantage of a transgenic fly line with an integrated bacterial artificial chromosome (BAC) containing the entire *hb* genomic locus that was modified to add a hemagglutinin (HA) epitope tag (Kohwi et al., 2013). The HA tag acts as a proxy for endogenous *hb* expression (Hb^HA^), and early-born neurons are identified by co-expression of Eve and HA (Hafer et al., 2022; Kohwi et al., 2013; Lucas et al., 2021).

Consistent with our previous studies (Kohwi et al., 2013), we found that misexpression of Dan^FL^ extended neuroblast early competence compared to misexpression of Hb alone (Hb alone: 5.4 Eve^+^HA^+^ neurons, N=8 embryos, 143 hemisegments, STD=0.88; Dan^FL^: 7.4 Eve^+^HA^+^ neurons, N=11 embryos, 184 hemisegments, STD=1.7, P value <0.0001). In contrast, misexpressing Dan^ΔPSQ^, in which the conserved 51 amino acid Psq sequence was deleted nearly abolished this phenotype (5.7 Eve^+^HA^+^ neurons, N=5 embryos, 90 hemisegments, STD=1.5, P value <0.0001) (Fig. 1B). By expressing each construct in salivary glands using a *forkhead*-gal4 driver (Henderson and Andrew, 2000), we confirmed that Dan^FL^, but not Dan^ΔPSQ^, binds DNA (Supp. Fig. 1Aii). Thus, the Psq DNA-binding domain is necessary for Dan function in neuroblast competence. In contrast, the Psq domain was not sufficient (5.3 Eve^+^HA^+^ neurons, N=5 embryos, 100 hemisegments, STD=1.3, P value=NS) (Supp. Fig. 1B).

Surprisingly, we found that misexpression of two copies of Dan^FL^ compared to just one copy did not increase competence extension. In fact, increasing Dan levels by the additional UAS transgene abolished competence extension entirely (5.1 Eve^+^HA^+^ neurons, N=3 embryos, 50 hemisegments, STD=2.1, P value=NS) (Fig. 1C). To control for the possibility that additional UAS transgenes dilutes the Gal4 protein, we replaced one of UAS-Dan transgenes for UAS-GFP, thus equalizing the total UAS binding sites in the animal, and the competence extension phenotype was fully restored (7.2 Eve^+^HA^+^ neurons, N=4 embryos, 62 hemisegments, STD=1.8, P value <0.0001) (Fig. 1C). This data indicate that Dan function is dose-sensitive. Using the prediction program IUPred3 (Erdos et al., 2021), we found that Dan protein is highly intrinsically disordered outside the Psq DNA-binding domain and is predicted to be 65% disordered (Fig. 1D). Studies have shown a high correlation between dose-sensitive proteins and high protein disorder (Vavouri et al., 2009), and these properties are frequently attributed to proteins that undergo liquid-liquid phase separation (LLPS) (Bolognesi et al., 2016; Martin and Mittag, 2018). Phase-separating proteins form condensates, or “hubs” within cells (Cho et al., 2018; Kent et al., 2020; Mir et al., 2017; Palacio and Taatjes, 2022; Sabari et al., 2018). Consistently, in fixed embryo preparations stained with an anti-Dan antibody, we found that Dan protein distributes into condensates of various sizes in addition to a more diffuse “haze” throughout the nucleoplasm (Fig. 1D).

### Dan has opposing DNA-binding and phase-separation functions

To examine Dan protein distribution in live cells, we transfected *Actin5C* (*Ac5*) promoter-driven constructs expressing GFP-fused Dan protein in Schneider 2 (S2) insect culture cells, a *Drosophila* embryo-derived cell line that does not express Dan endogenously (Cherbas et al., 2011). Dan:GFP distribution in S2 cells was similar to that of endogenous Dan in neuroblasts as well as *worniu-Gal4* driven (Albertson et al., 2004) expression of Dan:GFP live-imaged in dissociated neuroblasts, forming discrete foci among a diffuse “haze” (Fig. 2A). Using S2 cells, we expressed Dan:GFP harboring a glutamic acid-to-lysine (E45K) point mutation (Dan^E45K^:GFP) (Emerald et al., 2003; Tanaka et al., 2001; Trost et al., 2016; Wang et al., 1999) in the predicted DNA binding helix of the Psq domain, which has been shown previously to reduce, but not eliminate Dan function in antenna development (Emerald et al., 2003). We observed that the size of the condensates increased, and the general haze of uniform Dan protein distribution decreased. Surprisingly, we found that the size of the Dan condensates was proportional to the degree of impairment of the DNA-binding domain. We made increasingly severe DNA-binding domain deletions: in addition to the point mutation in the DNA-binding domain (Dan^E45K^:GFP), we also deleted of a core set of five amino acids in the same region as the E45K point mutation (Dan^ΔESTLR^:GFP), and finally deleted the entire Psq domain (Dan^ΔPSQ^:GFP). As the DNA-binding impairment increased, the size of the Dan condensates increased, while the number decreased (Fig. 2Aiii, Supp. Fig. 2Ai). As we did not find differences in the total GFP signal between the constructs, we conclude that impairing DNA-binding causes Dan protein to coalesce into steadily larger condensates (Supp. Fig. 2A). Consistent with liquid properties of the condensates, we found that both Dan^ΔE45K^:GFP and Dan^ΔPSQ^:GFP form highly circular foci (Supp. Fig. 2A) (Alberti et al., 2019; Strom et al., 2017; Zhu and Brangwynne, 2015).

**Figure. 2.**
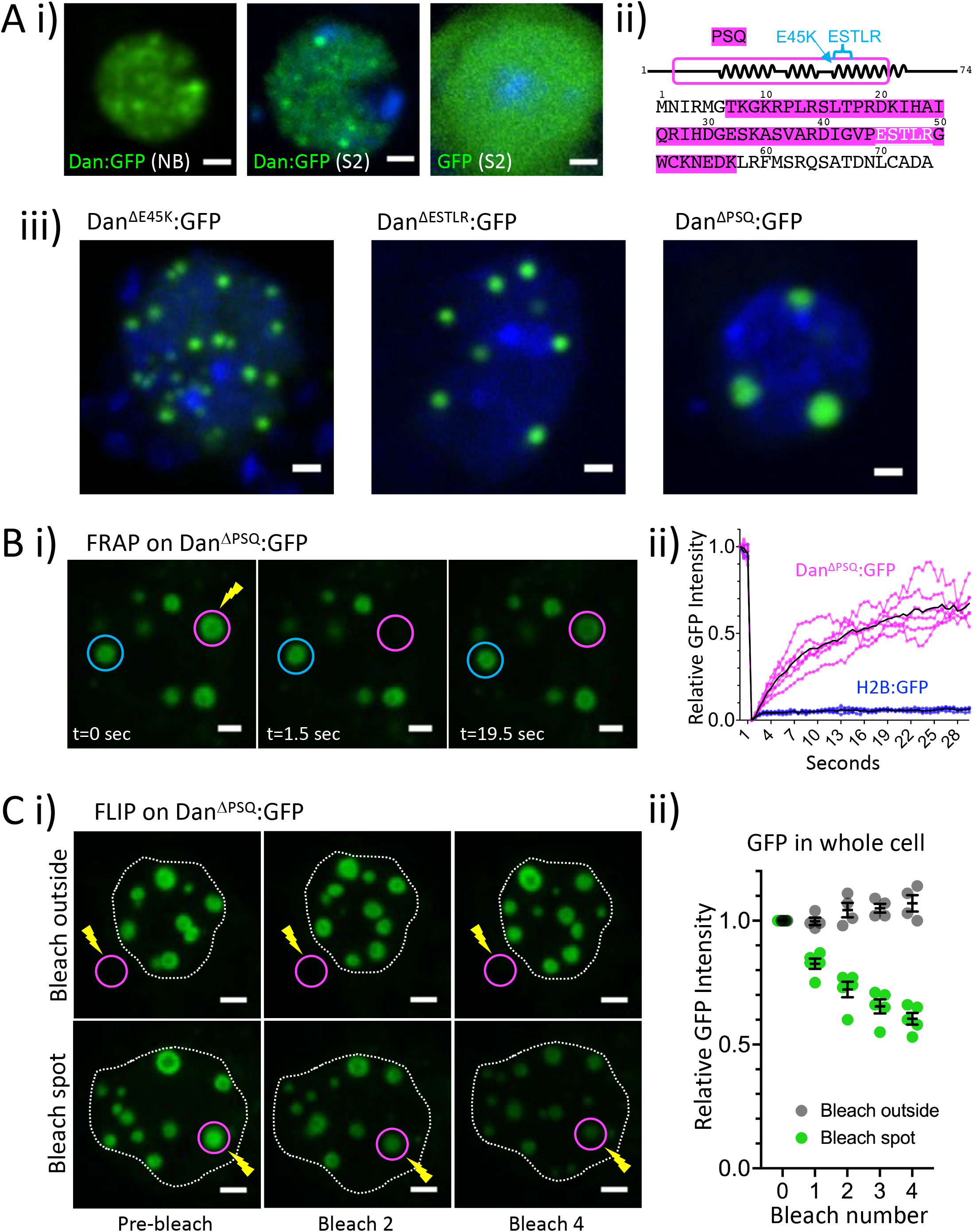
Dan has opposing DNA-binding and phase-separation functions. **A)** i, Dan:GFP expression in live dissociated neuroblast at stage 14 (NB), Dan:GFP or GFP alone in live S2 cells. S2 cells have Hoechst stain marking DNA. Scale bars, 1μm. **ii**, Schematic of the Dan N-terminus, showing the PSQ, ESTLR, and E45K mutations. **iii**, Expression of Dan DNA-binding mutants. Hoechst stain marks DNA. Scale bars, 1μm. **B) i**, Representative images of FRAP on a Dan^βPSQ^:GFP droplet in S2 cells 48hrs after transfection. Time points before bleach (t=0), immediately after bleach (t=1.5 sec), and after recovery (t=19.5 sec). Control (no bleach) droplet also shown. Scale bars, 1μm. **ii**, Quantification of signal recovery for bleached Dan^βPSQ^:GFP and H2B:GFP spots after FRAP. GFP relative intensity of “1” corresponds to average GFP signal of the first three images before bleaching. 5 cells were analyzed for each condition. **C) i**, Representative images of FLIP on Dan^βPSQ^:GFP in S2 cells 48hrs after transfection. Bleaching was repeated a total of 4 times, either on a single droplet or just outside of the nucleus. Scale bars, 1μm. **ii**, Quantification of total relative fluorescent signal from a z-projection through the entire S2 cell after FLIP bleaching. GFP relative intensity of “1” corresponds to average GFP signal of the first three images before bleaching. See also Supplemental Figure 2.

Together with the high intrinsic disorder and dose-sensitive properties, the data suggest that Dan is a phase-separating protein. To test this hypothesis further, we performed fluorescence recovery after photobleaching (FRAP). Photobleached LLPS droplets have been shown to recover their fluorescence on the scale of seconds to minutes due to rapid diffusion of protein in and out of droplets, while solid aggregates can take tens of minutes to hours to turn over (McSwiggen et al., 2019; Taylor et al., 2019). We found that Dan^ΔPSQ^:GFP foci undergo rapid signal recovery (Fig. 2B), indicating a rapid exchange of protein (Shapiro et al., 2021; Sprague and McNally, 2005; Taylor et al., 2019), in stark contrast to Histone2B:GFP (H2B:GFP), a stably bound protein that is commonly used as a FRAP control, which does not recover fluorescence in the time scale tested (Fig. 2B, Supp. Fig 2B) (Hansen et al., 2017; Mazza et al., 2012; Mir et al., 2018; Teves et al., 2016). Importantly, for proteins with liquid properties, FRAP recovery correlates with size of the LLPS hubs, as larger volume protein hubs require more diffusion and exchange between bleached and unbleached protein than in the smaller volume hubs (Sprague and McNally, 2005). We found that Dan condensate recovery times after FRAP were indeed proportional to size. The large droplets formed 48hrs after Dan^ΔPSQ^:GFP transfection recovered to 50% fluorescence intensity within an average of 11.5 seconds, whereas the small droplets formed 24hrs after Dan^ΔE45K^:GFP transfection recovered to 50% fluorescence intensity within an average of 4.8 seconds (P value < 0.001, N=5-6 cells) (Supp. Fig. 2B). In all cases, a control region or droplet distal to the bleached droplet was largely unaffected by the immediate bleach event (Supp. Fig. 2B).

Interestingly, we found that repeated photobleaching of a single Dan^ΔPSQ^:GFP droplet, or FLIP (fluorescence loss in photobleaching) (Chen and Huang, 2001; Phair and Misteli, 2000; White and Stelzer, 1999), led not only to an immediate loss of fluorescence in the bleached droplet, but a gradual loss of fluorescence in all of the droplets in the cell (Fig. 2C). We repeatedly photobleached a single droplet four times and imaged the whole nucleus three-dimensionally between each bleach event. We used the same bleach setting (area, laser intensity, and duration), and for control nuclei, we bleached just outside the nucleus. In control nuclei, the total fluorescence intensity remained the same, or increased slightly over the four bleach cycles. In contrast, repeated bleaching of a single droplet resulted in a significant reduction in total fluorescent intensity over time (Fig. 2C). We note that each bleach event only affected the targeted droplet and had no immediate effect on neighboring droplets within a five second window, but after 30 seconds, the fluorescent intensity of neighboring droplets decreased (Supp. Fig. 2C). Thus, while Dan^ΔPSQ^:GFP hubs appear as stable, independent units, they are highly interconnected, and individual Dan protein molecules move freely and rapidly between them. Together, we conclude that Dan has DNA-binding and LLPS properties, which exert opposing effects on Dan protein distribution within nuclei.

### Dan has a LLPS domain that is necessary for competence regulation

A recent study developed a protein prediction tool to identify LARKS domains (low-complexity, amyloid-like, reversible, kinked segment), which are weakly self-interacting and are frequently found in phase-separating proteins and membrane-less organelles (Hughes et al., 2021; Hughes et al., 2018). The kinked structure of LARKS domains facilitates reversible binding, which allows for dynamic protein organization. We found a predicted LARKS domain near the Dan Psq motif (Fig. 3A). To test the role of the LARKS domain on Dan function *in vivo*, we generated a myc-tagged UAS-Dan^ΔLARKS^ transgenic fly, in which 13 amino acids of the predicted LARKS domain, primarily glycines, were deleted (Fig. 3A). Importantly, in salivary gland cell polytene chromosomes, Dan^ΔLARKS^ still binds widely across the genome, indicating that the LARKS domain is independent of the DNA-binding domain, and DNA-binding is not impaired in Dan^ΔLARKS^ (Fig. 3B). However, neuroblast competence extension was abrogated upon misexpression of Dan^ΔLARKS^ (Dan^FL^: 7.3 Eve^+^HA^+^ neurons, N=5 embryos, 94 hemisegments, STD=1.4; Dan^ΔLARKS^: 5.8 Eve^+^HA^+^ neurons, N=6 embryos, 100 hemisegments, STD=1.4, P value <0.0001) (Fig. 3C). We conclude that the LARKS domain is necessary for Dan function in neuroblast competence *in vivo*.

**Figure 3.**
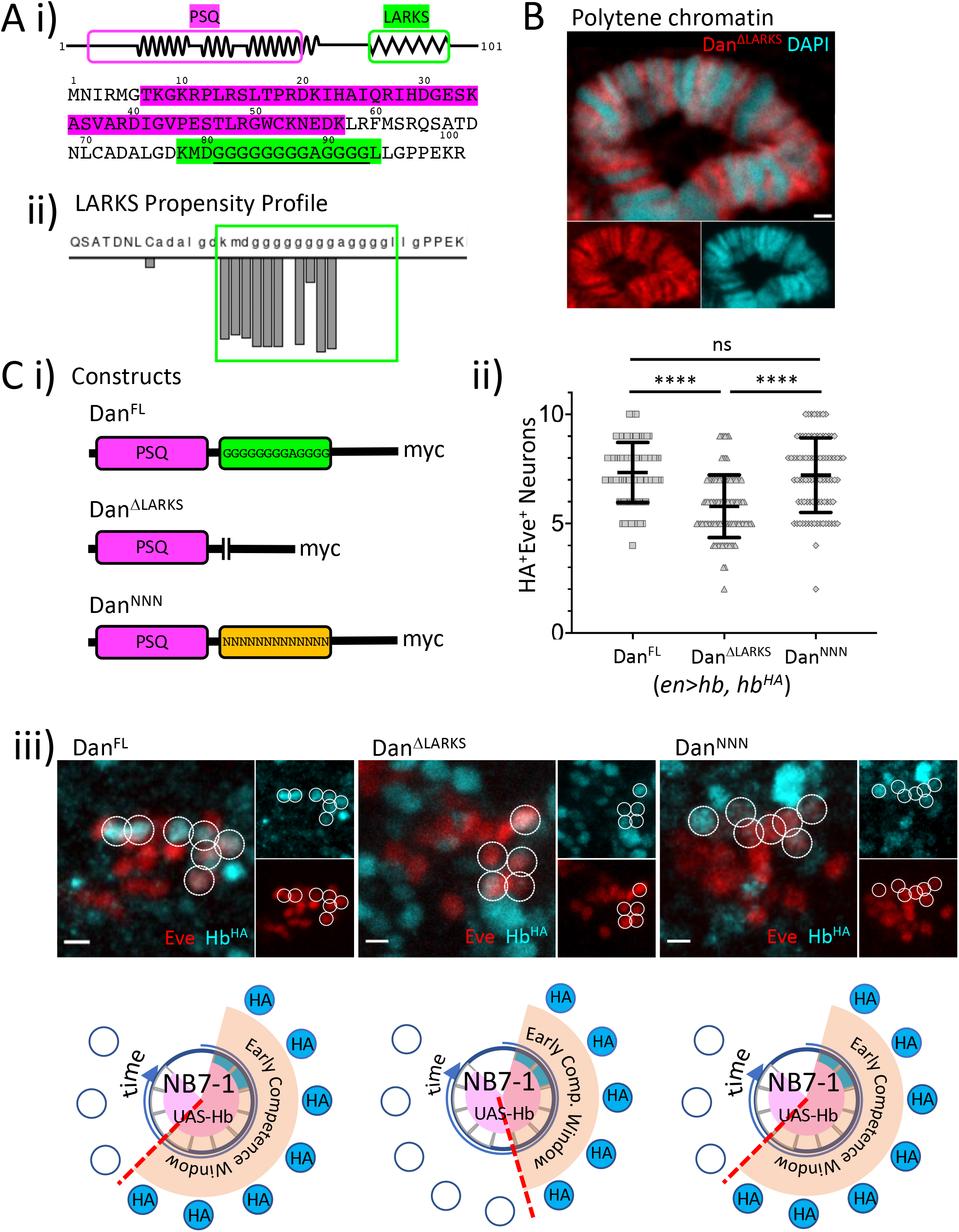
Dan has a LLPS domain that is necessary for competence regulation. **A) i**, Schematic diagram of Dan protein showing the Psq motif DNA-binding domain (magenta) and the LARKS domains (green). Amino acid sequence shown below diagram. Underlined region indicates the animo acids deleted in the Dan^βLARKS^ construct. **ii**, LARKS propensity profile identifies the glycine-rich region predicted to be the LARKS domain. **B)** In polytene chromosomes, Dan^βLARKS^, detected by immunostaining for the myc tag, can still bind DNA. **C) i**, Schematic diagram of structure function mutant constructs. **ii-iii**, Upon loss of LARKS domain, Dan’s ability to extend neuroblast competence is abrogated. This function is fully rescued when Dan’s LARKS domain is replaced with a poly-asparagine stretch (Dan^NNN^). Images are of single representative hemisegments of the data shown in ii. Each data point represents a single hemisegment, and mean, standard deviation, and t-test significance is shown.

To determine whether the loss of competence extension upon misexpression of Dan^ΔLARKS^ in neuroblasts is due to loss of Dan’s LLPS properties, we performed a rescue experiment by replacing the deleted Dan LARKS domain with a poly-asparagine repeat (Dan^NNN^). Asparagines are found in high abundance in prions and prion-like domains, which are known to phase-separate (Du, 2011; Lu and Murphy, 2015; Michelitsch and Weissman, 2000; Molliex et al., 2015). Poly-asparagine repeats are common in invertebrates, and particularly in *Drosophila* transcription factors, suggesting that such repeats may be functional in nuclear activity (Faux et al., 2005). We found that the poly-asparagine LARKS domain replacement (Dan^NNN^) was sufficient to rescue the Dan LARKS domain deletion (7.2 Eve^+^HA^+^ neurons, N=5 embryos, 89 hemisegments, STD=1.7, P value <0.0001), and was able to extend early competence to the same extent as Dan^FL^ (Fig. 3C). We conclude that the phase separation function of the Dan LARKS domain is conserved, and is necessary for its function in neuroblast competence regulation.

### Dan interacts with nucleopore complex proteins, which are required for terminating neuroblast early competence

One unexpected outcome of Dan^ΔPSQ^:GFP distribution in S2 cells was that in addition to forming large protein hubs, the droplets became relocalized to the nuclear periphery. Measurements of pixel intensity through a 3D projection of the cells shows that the droplet average peak pixel intensity lies near the edge of the nucleus (N=3 cells, 20 droplets), while Dan^FL^:GFP foci were distributed throughout the nucleus (N=4 cells, 20 measurements) (Fig. 4A). Thus, we hypothesized that when Dan is free from genome-binding, the protein defaults to preferentially interacting with itself and with proteins at the nuclear periphery. To identify Dan-interacting proteins in neuroblasts, we expressed myc-tagged Dan specifically in neuroblasts with *worniu-Gal4* driver, and performed immunoprecipitation-mass spectrometry (IP-MS) using anti-myc beads (Fig. 4B). Consistent with Dan’s propensity to localize to the nuclear periphery, Gene Ontology analysis of significantly enriched proteins identified in the IP-MS revealed a high prevalence for association with the nuclear periphery and the nuclear pore complex (NPC), and we identified multiple NPC subunits among the enriched proteins (Fig. 4B, Supp. Fig. 4A). NPC proteins were also enriched among the immunoprecipitated lysate in wild type embryos immunoprecipitated with an anti-Dan antibody, indicating a strong association with proteins of the NPC (Supp. Fig. 4B).

**Figure 4.**
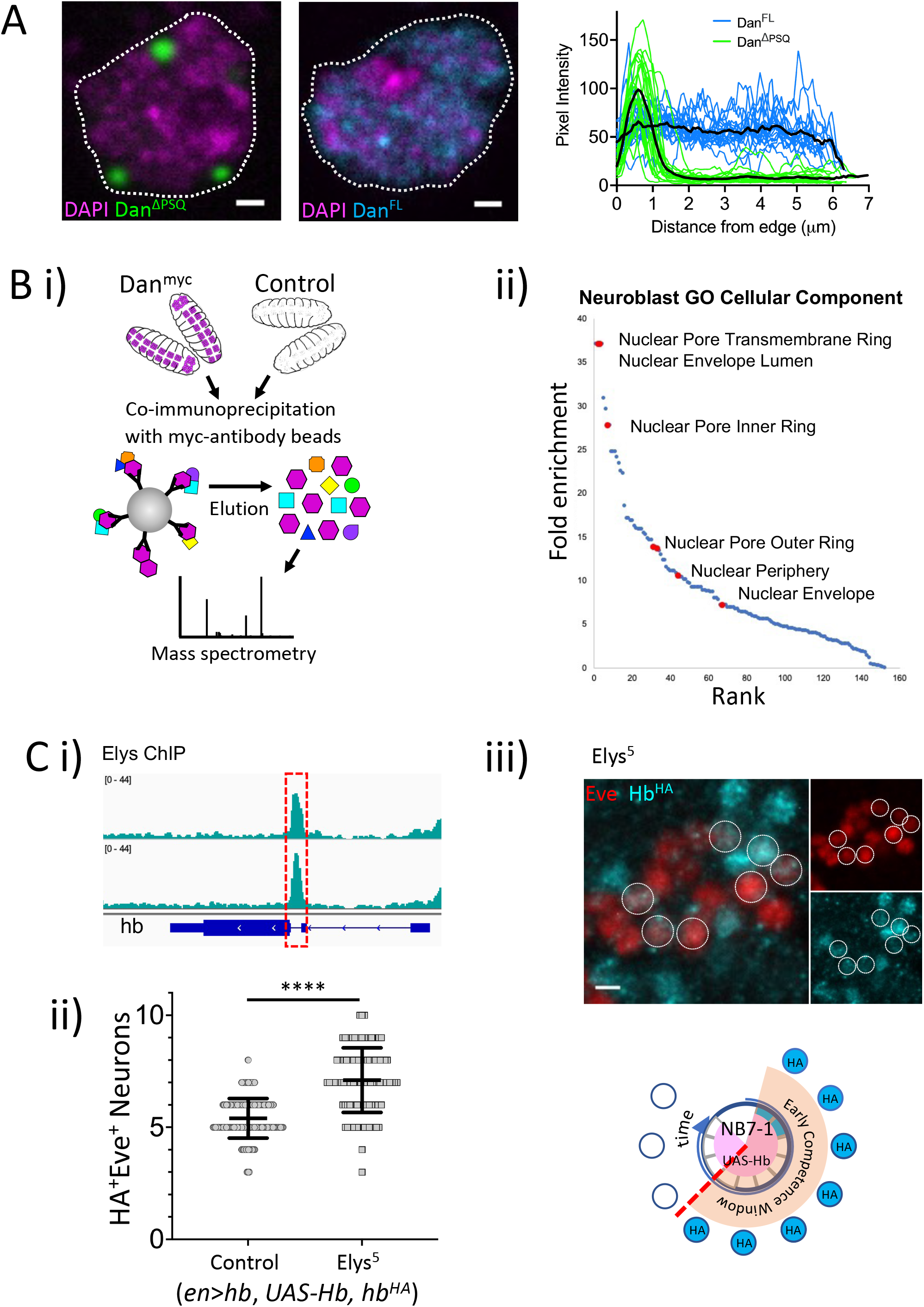
Dan interacts with nucleopore complex proteins, which are required for terminating neuroblast early competence. **A)** Upon deletion of the DNA-binding domain, Dan protein hubs relocalize to the nuclear periphery. Graph shows GFP signal intensity plotted as distance from edge of nucleus comparing Dan^FL^ and Dan^βPSQ^. **B)** Schematic diagram of IP-MS. Briefly, lysates from embryos expressing myc-tagged Dan were immunoprecipiated with anti-myc antibodies, and subjected to mass spectrometry. Embryos without any myc tags were used as controls. Gene ontology of cellular component revealed high enrichment of proteins associated with the nuclear pore complex (NPC). **C) i**, Analysis of published ChIP-seq datasets for Elys show Elys binds to the *hb* intron, recently identified to be necessary and sufficient for gene relocation to the nuclear lamina (see text for details). **ii**, In loss of function Elys mutants, the early competence window is prolonged. **iii**, image is of a single representative hemisegment of the data shown in ii. Each data point represents a single hemisegment, and mean, standard deviation, and t-test significance is shown. See also Supplemental Figure 3.

Elys, one of the NPC proteins found in the IP-MS analyses, is a core scaffold protein of the NPC and has been shown to play a role in genome architecture regulation (Shevelyov, 2020). Elys binds across the genome at sites that intersect with multiple other NPC proteins (Gozalo et al., 2020; Pascual-Garcia et al., 2017; Shevelyov, 2020) and has also been shown to bind and relocate specific loci to the NPC (Scholz et al., 2019). Taking advantage of published Elys chromatin immunoprecipitation with sequencing (ChIP-seq) datasets from third instar *Drosophila* larvae brains, we found that Elys binds specifically at the *hb* gene intron (Fig. 4C) (Pascual-Garcia et al., 2017). We recently identified the *hb* intronic element (IE) to be necessary and sufficient for gene relocation to the nuclear periphery and termination of the NB7-1 early competence window (Lucas et al., 2021). Using our competence assay in loss-of-function Elys mutants (Elys^5^) (Hirai et al., 2018), we found an increase in early-born neurons in the Elys^5^ mutant background (7.1 Eve^+^HA^+^ neurons, N=5 embryos, 106 hemisegments, STD=1.4, P value <0.0001) (Fig. 4C). Together, these data indicate that Dan functions through balancing its interaction with the genome, itself, and other nuclear architecture regulators, such as the NPC, supporting a model by which phase-separation plays an important role in nuclear architecture reorganization during neuroblast competence transitions (Fig. 5).

**Figure 5.** Model for Dan function in neuroblasts. **A)** Dan protein distribution is influenced by the opposing forces of two protein properties, DNA-binding and self-association. When the DNA-binding function is impaired, Dan protein coalesces into large, liquid hubs that relocalize to the nuclear periphery. IP-MS shows Dan strongly interacts with proteins of the NPC. **B)** In the embryo, neuroblasts express high levels of Dan initially, and rapidly downregulate Dan at mid-embryogenesis at which time the *hb* gene relocates to the nuclear lamina for heritable silencing. Our results support a model in which loss of Dan protein releases target genes, which can subsequently become subject to nuclear architecture reorganization.

## Discussion

Dan is a member of a large superfamily of evolutionarily conserved proteins characterized by a pipsqueak (Psq) DNA-binding domain and is closely related to the CENP-B/Transposase family (Siegmund and Lehmann, 2002). Dan is robustly expressed by all newly born neuroblasts, but is transiently downregulated at mid-embryogenesis, and this downregulation is required for the early-born neuron gene *hb* to relocate to the neuroblast nuclear lamina and terminate the early competence window (Kohwi et al., 2013). Thus, the absence of Dan is required for nuclear architecture reorganization. Misexpressing Dan in neuroblasts to override this transient downregulation blocks *hb* gene-lamina relocation and extends the early competence window, suggesting that Dan acts to stabilize the existing genome organization. Increasing studies have identified phase-separating properties in nuclear factors, revealing an important role for LLPS in subnuclear organization. In addition to formation of the nucleolus, phase-separation has been implicated in heterochromatin compaction (Larson et al., 2017; Liu et al., 2020; Sanulli et al., 2019; Strom et al., 2017), transcriptional regulation (Cho et al., 2018; Sabari et al., 2018), and formation of topologically associated domains (Wang et al., 2021). In this study, we found that Dan is a phase-separating protein, forming large hubs when DNA-binding is impaired. These hubs are interconnected and characterized by rapid protein exchange, suggesting high protein mobility and strong self-interactions. Thus, Dan’s propensity to bind the genome through its Psq motif DNA-binding domain and its propensity to self-associate act in a push-pull relationship on Dan protein distribution within nuclei. One can easily imagine how such opposing properties within the same protein could exert mechanical forces to organize or stabilize their local genomic environment, perhaps through dynamic restructuring (Shin et al., 2018). An attractive possibility is that Dan binds and gathers multiple genomic loci in local phase-separated hubs, and at mid-embryogenesis, neuroblasts transiently downregulate Dan, releasing the target loci, thereby allowing a new genome organization to take place.

Deletion of the DNA-binding domain resulted not only in the formation of large LLPS hubs, but these hubs relocalized to the nuclear periphery. Consistently, IP-MS revealed strong interactions with subunits of the NPC, including Elys, a core scaffold protein of the NPC, which has a DNA-binding domain (Shevelyov, 2020). Our analysis of published ChIP-seq datasets showed Elys binding at the *hb* intron, and loss of Elys extended neuroblast early competence *in vivo*. We recently found that the *hb* intron is a gene mobility element, both necessary and sufficient for gene relocation to the nuclear lamina, and is targeted by the Polycomb (PcG) chromatin factors (Lucas et al., 2021). There is increasing evidence that NPC proteins and PcG proteins work together in gene regulation (Gozalo et al., 2020; Jacinto et al., 2015), and Pipsqueak, the namesake of the protein superfamily, has been shown to recruit PcG proteins to PcG response elements (Gutierrez-Perez et al., 2019; Huang et al., 2002; Schwendemann and Lehmann, 2002). Interestingly, while Dan was found to interact with multiple NPC proteins, including Elys, Dan does not interact with PcG proteins (Supp. Fig. 4). Future studies will address how Dan interacts with other nuclear architecture regulators and the role of phase-separation in these contexts.

## Materials and Methods

### Fly lines

Wild-type *(w1118), Psc*^*e24*^(BDSC#24155), *1407-GAL4* (BDSC#8751), *UAS-GFP* (BDSC#32184), *wor-GAL4* (BDSC#56553), *fkh-GAL4* (BDSC#78060), *GAL80*^*ts*^ (BDSC#7019), *en-GAL4* (chrom II, BDSC#46438), *UAS-hb* (Wimmer et al., 2000), *hb*^*HA*^ (Kohwi et al., 2013), *Elys*^*5*^ (Hirai et al., 2018) (kind gift from Dr. Kyoichi Sawamura, Tsukuba University, Japan), UAS-Dan^FL^ (this paper), UAS-Dan^GFP^ (this paper), UAS-Dan^Myc^ (this paper, called Dan^FL^ in Fig 3), *UAS-Dan*^*ΔPSQ*^ (this paper), *UAS-Dan*^*ΔPSQ-Myc*^ (this paper), *UAS-PSQ:GFP* (this paper), *UAS-Dan*^*ΔLARKS*^ (this paper), *UAS-Dan*^*NNN*^ (this paper). Transgenic flies were generated through PhiC31-mediated insertion at either attP2 or attP40. Flies were raised on a standard cornmeal/molasses medium at 25°C.

### Plasmids

UAS-Dan^FL^ was made by restriction cloning from Dan cDNA LD40883 and UAS plasmid PRVV70 (kind gift from Dr. Richard Mann, Columbia University). PRVV70 has a phi-C31 integrase binding site for genomic integration, a UAS/Gal4 binding site, and a multiple cloning site containing XbaI and KpnI restriction sites. The Dan 5’UTR and coding region were extracted by PCR from LD40883 (#5984 DRGC), and XbaI and KpnI restriction sites were added. Ac5-Dan^FL^ was made by DNA assembly using NEBuilder HiFi DNA Assembly Cloning Kit (NEB #E5520S) from pAc5.1C-FLuc-V5His6 (Addgene plasmid #21183), Ac5-STABLE1-neo (Addgene plasmid #32425) and UAS-Dan^FL^. All other plasmids were generated by DNA assembly or site-directed mutagenesis using UAS-Dan^FL^ or Ac5-Dan^FL^ as the backbone. All plasmid sequences from this study can be downloaded from https://benchling.com/gtb2109/f_/HnCjmTJr-dan-paper/.

### S2 cells

S2 cells were grown in T25 flasks (Thermo Scientific, #156367) at 25°C in Schneider’s Drosophila Medium (Gibco #21720024) with 10% heat inactivated FBS, and cultured following standard protocols (Luhur et al., 2019). For transfection, 1×10^6^ cells were seeded onto 35mm glass bottom dish (Cellvis #D35-20-1.5-N). After 24 hours, cells were transfected with 500ng plasmid DNA and a 1:50 Effectene transfection reagent ratio (Qiagen #301425).

### Immunohistochemistry

Embryos from overnight collection at 29°C were immunostained following standard protocols (Lucas et al., 2021; Rothwell, 2000). Briefly, embryos were fixed in a 1:1 mixture of 4% formaldehyde in PEM buffer (0.1M Pipes, 1mM MgS04, 2mM EGTA) and N-heptane and rocked for 22 min at room temperature. Embryos were devitellinized by vigorous shaking in a 1:1 mixture of methanol:heptane and washed with PBS-0.1% Tween 20 (PBST) before staining. Embryos were incubated in primary antibodies diluted in PBST overnight at 4°C, secondary antibodies at room temperature for 1.5 hours, and streptavidin for 20 mins at room temperature.

Salivary glands were fixed and immunostained following standard protocol with minor modifications (Kuhn et al., 2020). Briefly, salivary glands were dissected from L3 larvae grown at 16°C after a 4-hour heat induction at 29°C. Dissected glands were fixed for 90 seconds in 2% PFA/45% Acetic Acid, then transferred to a droplet of 45% Acetic acid on a siliconized coverslip. The glands were squashed, flash frozen in liquid nitrogen, then washed in PBST. Squashes were incubated in antibodies diluted in PBST overnight at 4°C, secondary antibodies at room temperature for 1 hour, and DAPI for 2 mins at room temperature.

### Photobleaching assays

#### Fluorescence Recovery After Photobleaching (FRAP)

Cells were imaged every 500ms for 60 frames on a Zeiss LSM 700 Confocal with a 63x objective with Immersol 518F immersion oil (Carl Zeiss #444960). After the third image, a circular region with a diameter ∼1μm was bleached using the 488nm laser at 100% power for 54.03μs, after which imaging continued for the remaining frames. Recovery was measured as fluorescence intensity of the photobleached or control area normalized to the intensity of the bleached spot immediately after bleaching.

#### Fluorescence Loss In Photobleaching (FLIP)

A z-stack of a transfected cell was taken prior to bleaching, after which the cell was imaged every 500ms for 10 frames, and bleached using the same conditions as for FRAP. After the 10 frames, 3D imaging of the entire nucleus was repeated. 3D imaging was performed a total of 5 times, and bleaching a total of 4 times. Bleaching was repeated on the same aggregate or in the same spot outside the nucleus. Fluorescence loss was measured in either the whole cell, by measuring the relative fluorescence intensity of combined Z-stack over time, or in individual droplets by measuring the relative fluorescence intensity of individual droplets in a single cell over time. Recovery of the bleached spot immediately before and after bleaching was measured as fluorescence intensity of the photobleached or control spots normalized to the intensity of the bleached spot immediately after bleaching.

### Antibodies used

Primary antibodies: anti-Eve (Mouse, #3C10 Developmental Studies Hybridoma Bank), anti-HA (Rat, #11867423001 Sigma Roche), anti-Dan (Rabbit) (Kohwi et al., 2013)), anti-Myc (Rabbit, #ab9106 Abcam), anti-Psc (Mouse, #6E8 Developmental Studies Hybridoma Bank). Secondary antibodies: Goat anti-Rabbit IgG, Alexa Fluor 555 (#A21429 Invitrogen), Donkey anti-Rat IgG, Biotin-SP (#712065153 Jackson ImmunoResearch), Strepavidin Cy3 (#SA1010 Invitrogen) Donkey anti-Mouse IgG, Alexa Fluor 647 (#715605151 Jackson ImmunoResearch), Goat anti-Mouse IgG, DyLight 550 (#A90516D3 Bethyl Laboratories), Goat anti-Rabbit IgG, Alexa Fluor 488 (#A11034 Invitrogen). DAPI (#D3571 Invitrogen). S2 cells were stained with NucBlue (#R37605, ThermoFisher).

### Neuroblast dissociation

UAS-Dan^GFP^, *wor*-gal4 embryos were collected after 3 hours of egg-laying at 25°C, and then incubated another 4 hours at 25°C until embryos were between 4-7 hours (embryonic stage 9-11). Embryos were dechorionated in 4% bleach for 3 mins, then rinsed for 2 mins in RO water. Dechorionated embryos were incubated in 95% ethanol for 5 mins, then washed in Chan and Gehring’s Balanced Saline (C&G) (Chan and Gehring, 1971). Embryos were homogenized in C&G-2%FBS, and cell suspension was plated on a round coverslip. Plated cells were incubated at 25°C for an additional 3 hours (∼ embryonic stage 14), then imaged.

### Co-Immunoprecipitation

Embryos were collected after 3 hours of egg-laying at 25°C, and then incubated another 4 hours at 25°C until embryos were between 4-7 hours (embryonic stage 9-11), during which Dan is primarily expressed in the nervous system (Kohwi et al., 2011). Protein was prepared using the Pierce™ c-Myc-Tag Magnetic IP/Co-IP Kit (ThermoScientific #88844), or Pierce™ MS-Compatible Magnetic IP Kit, protein A/G (ThermoScientific #90409) using Dan antibody or IgG, and bead-immobilized proteins were analyzed by mass-spectrometry.

### Mass Spectrometry

Mass Spectometry was performed at Columbia’s proteomics core directed by Dr. Rajesh Soni.

#### On-bead digestion of samples for mass spectrometry

Proteins bound to magnetic beads were washed five times with 200 µl of 50 mM ammonium bicarbonate and subjected to disulfide bond reduction with 5 mM TECP (room temperature, 30 min) and alkylation with 10 mM iodoacetamide (room temperature, 30 min in the dark). Excess iodoacetamide was quenched with 5 mM DTT (room temperature, 15 min). Proteins bound on beads were digested overnight at 37°C with 1 µg of trypsin/LysC mix. The next day, digested peptides were collected in a new microfuge tube and digestion was stopped by the addition of 1% TFA (final v/v), and centrifuged at 14,000 g for 10 min at room temperature. Cleared digested peptides were desalted on SDB-RP Stage-Tip and dried in a speed-vac. Peptides were dissolved in 3% acetonitrile/0.1% formic acid.

#### Liquid chromatography with tandem mass spectrometry (LC-MS/MS)

Desalted peptides were injected in an EASY-SprayTM PepMapTM RSLC C18 50cm X 75cm ID column (Thermo Scientific) connected to an Orbitrap FusionTM TribridTM (Thermo Scientific). Peptides elution and separation were achieved at a non-linear flow rate of 250 nl/min using a gradient of 5%-30% of buffer B (0.1% (v/v) formic acid, 100% acetonitrile) for 110 minutes with a temperature of the column maintained at 50 °C during the entire experiment. The Thermo Scientific Orbitrap Fusion Tribrid mass spectrometer was used for peptide tandem mass spectroscopy (MS/MS). Survey scans of peptide precursors are performed from 350 to 1500 m/z at 120 K full width at half maximum (FWHM) resolution (at 200 m/z) with a 2 × 10^5^ ion count target and a maximum injection time of 60 ms. The instrument was set to run in top speed mode with 3-second cycles for the survey and the MS/MS scans. After a survey scan, MS/MS was performed on the most abundant precursors, i.e., those exhibiting a charge state from 2 to 6 of greater than 5 × 10^4^ intensity, by isolating them in the quadrupole at 1.6 Th. We used higher-energy collision dissociation (HCD) with 30% collision energy and detected the resulting fragments in the ion trap. The automatic gain control (AGC) target for MS/MS was set to 5 × 10^4^ and the maximum injection time was limited to 30 ms. The dynamic exclusion was set to 30 s with a 10-ppm mass tolerance around the precursor and its isotopes. Monoisotopic precursor selection was enabled.

#### LC-MS/MS data analysis

Raw mass spectrometric data were analyzed using the MaxQuant environment v.1.6.1.0 (Cox and Mann, 2008) and Andromeda for database searches (Cox et al., 2011) at default settings with a few modifications. The default is used for first search tolerance and main search tolerance (20 ppm and 6 ppm, respectively). MaxQuant was set up to search with the reference Drosophila *melanogaster* proteome database downloaded from UniProt. MaxQuant performed the search trypsin digestion with up to 2 missed cleavages. Peptide and protein false discovery rates (FDR) were all set to 1%. The following modifications were used for protein identification and quantification: oxidation of methionine (M), acetylation of the protein N-terminus, and deamination for asparagine or glutamine (NQ). Results obtained from MaxQuant were used further analysis.

### Imaging and analyses

Imaging of embryo competence and salivary glands was done using a Zeiss LSM 700 Confocal microscope. Imaging of S2 cells and dissociated embryonic cells was done using a W1-Yokogawa Spinning Disk Confocal, 48hrs after transfection and incubation at 25°C. All image analyses were done using FIJI (Schindelin et al., 2012).

#### S2 cell protein distribution

Images were processed using FIJI by adding a Gaussian Blur filter, and 3D-Projection with Brightest Point Projection and Y-Axis rotation settings. A straight-line ROI was drawn from one edge of the cell to the other, passing through at least one unique droplet and the midpoint of the cell. A minimum of 5 straight-line ROIs were drawn per cell, and intensity of the line was measured using Plot Profile function. Measurements were normalized to background intensity outside of the nucleus.

#### S2 cell droplet comparisons

Droplets from each cell were manually thresholded and saved as ROIs. The number of droplets per cell was counted manually. Area and Circularity were measured using FIJI Measurements. Total pixel intensity was calculated by summing the z-stack and measuring mean gray value using FIJI Measurements (Schindelin et al., 2012).

### Other analyses

*Statistics:* We applied standard t-tests using Prism v8. Statistical significance was classified as follows: ^∗^ < 0.05, ^∗∗^ < 0.01, ^∗∗∗^ < 0.001, ^∗∗∗∗^ < 0.0001. Graphs show each data point used for statistical analysis, and bars indicate mean and standard deviation. Gene ontology (GO) analysis was performed using PatherDB (Mi et al., 2021). Dan intrinsic disorder was predicted using IUPred3, using IUPred3 long disorder analysis and medium smoothing (Erdos et al., 2021).

## Acknowledgements

We thank members of the Kohwi Lab for scientific discussion and feedback. We thank Dr. Natalia Molotkova and Ms. Sofiya Patra for technical assistance with experiments. We also thank Drs. Daniel Kalderon, Stavros Lomvardas, Richard Mann, and Tanguy Lucas for scientific discussion and feedback.

## Author contributions

G.B. and M.K. designed and executed the experiments, analyzed data and wrote the manuscript. M.J.L., R.J.C., and Y.H. assisted with experimental execution and data analysis. M.K. received support from NIH grants R00HD072035 and R01HD092381, Rita Allen Foundation, and Whitehall Foundation. G.B. received support from the NIH T32 training grant GM879818.

