## Supplemental Figures for "Nuclear architecture protein Distal antenna balances genome-binding and phase-separation properties to regulate neuroblast competence"

Supplemental Figure legends

<u>Supplemental Figure 1 (Supplement to Figure 1)</u>
**Ai:** Schematic of UAS constructs inserted into identical genomic sites by phi-C31 integration.
**Aii:** Squashed salivary glands immunostained for myc tag fused to $Dan^{FL}$ and $Dan^{\Delta PSQ}$. Scale bars, 10µm.
**Bi:** NB7-1 lineage early competence assay comparing control (misexpressing Hb only) and PSQ:GFP. Scale bars, 3µm.
**Bii:** Quantification of data shown in i. Each data point represents a single hemisegment, and mean, standard deviation, and t-test significance is shown.

<u>Supplemental Figure 2 (Supplement to Figure 2)</u>

**Ai:** Quantification of the number of droplets per cell for $Dan^{\Delta E45K}$:GFP and $Dan^{\Delta PSQ}$:GFP. 4-5 cells were analyzed for each condition.
**Aii:** Quantification of the largest 2D area in $\mu m^2$ for individual $Dan^{\Delta E45K}$:GFP and $Dan^{\Delta PSQ}$:GFP droplets. 4-5 cells were analyzed for each condition, with 115 total droplets analyzed for $Dan^{\Delta E45K}$:GFP and 48 droplets analyzed for $Dan^{\Delta PSQ}$:GFP.
**Aiii:** Quantification of total fluorescent intensity from a z-projection of $Dan^{\Delta E45K}$:GFP and $Dan^{\Delta PSQ}$:GFP expressing cells. 4-5 cells were analyzed for each condition.
**Aiv:** Quantification of circularity for individual $Dan^{\Delta E45K}$:GFP and $Dan^{\Delta PSQ}$:GFP droplets. 4-5 cells were analyzed for each condition, with 115 total droplets analyzed for $Dan^{\Delta E45K}$:GFP and 48 droplets analyzed for $Dan^{\Delta PSQ}$:GFP. See methods for details.
**Bi:** Representative images of FRAP on a $Dan^{\Delta E45K}$:GFP droplet in S2 cells 24hrs after transfection and H2B:GFP region in S2 cells 24hrs after transfection. Time points before bleach (t=0), immediately after bleach (t=1.5 sec), and after recovery (t=19.5 sec). Control (no bleach) droplet or regions also shown. Scale bars, 1µm.
**Bii:** Quantification of signal recovery for bleached $Dan^{\Delta E45K}$:GFP and H2B:GFP spots after FRAP. 5 cells were analyzed for each condition.
**Biii:** Quantification of non-bleached spots for $Dan^{\Delta PSQ}$:GFP, H2B:GFP after 24 or 48 hours, and $Dan^{\Delta E45K}$:GFP
**C:** Quantification of fluorescent intensity of droplets from a single, representative $Dan^{\Delta PSQ}$:GFP cell. Pink traces represent the bleached droplet, and black traces represent non-bleached droplets in the same cell.

<u>Supplemental Figure 3 (Supplement to Figure 4)</u>
**A** Ranking of Gene Ontology (GO) Cellular Component Analysis categories by fold-change in both Dan antibody IP-MS. Nuclear periphery categories are highlighted.
**B** Ranking of identified proteins by fold-change in both Dan antibody IP-MS and Dan-myc IP-MS. Dan, lamin, and nuclear pore complex proteins are highlighted.

Supplemental FIGURE 1

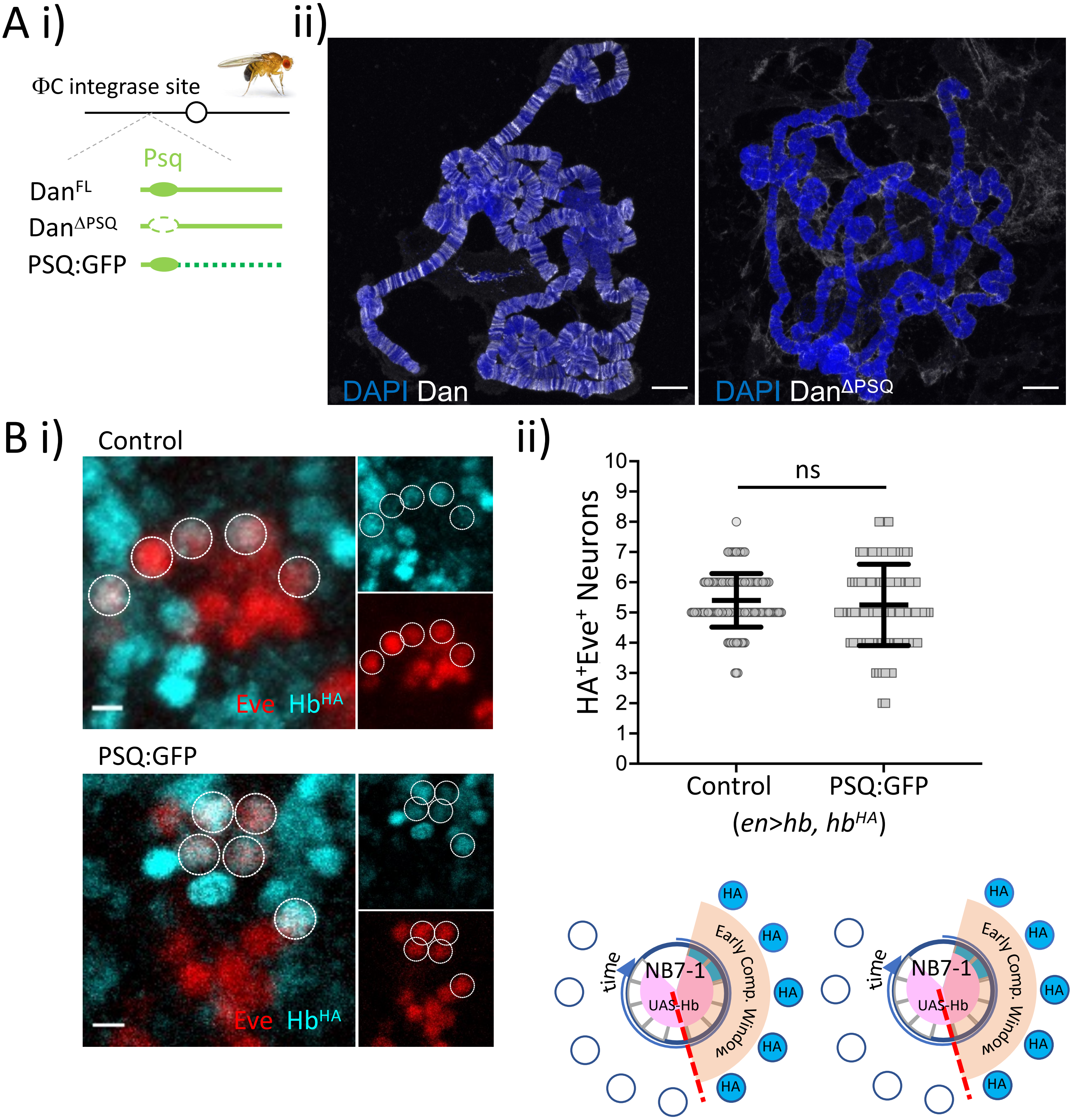

Supplemental FIGURE 2

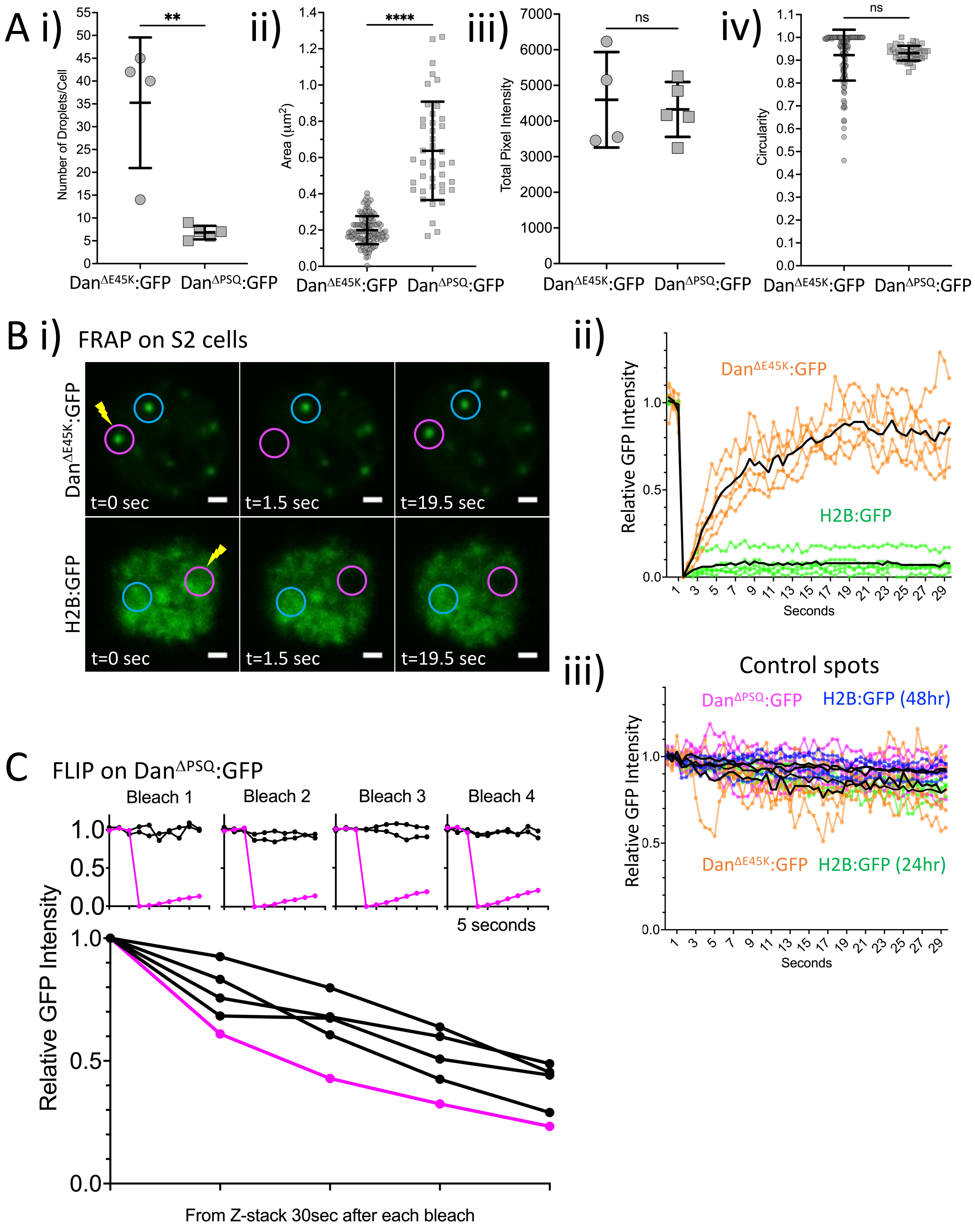

Supplemental FIGURE 3

A

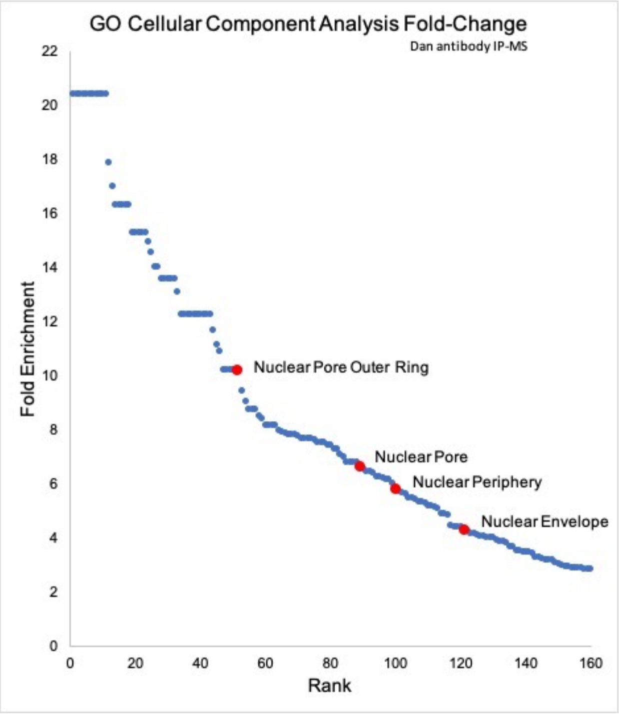


B

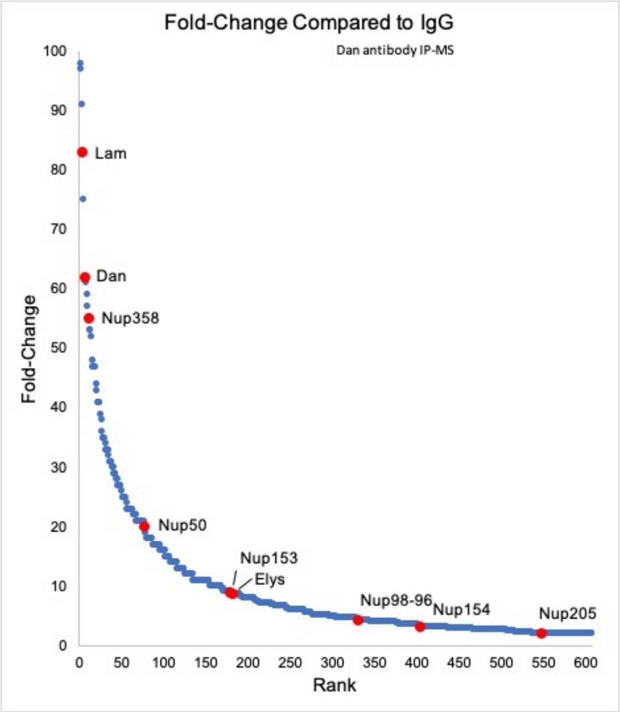


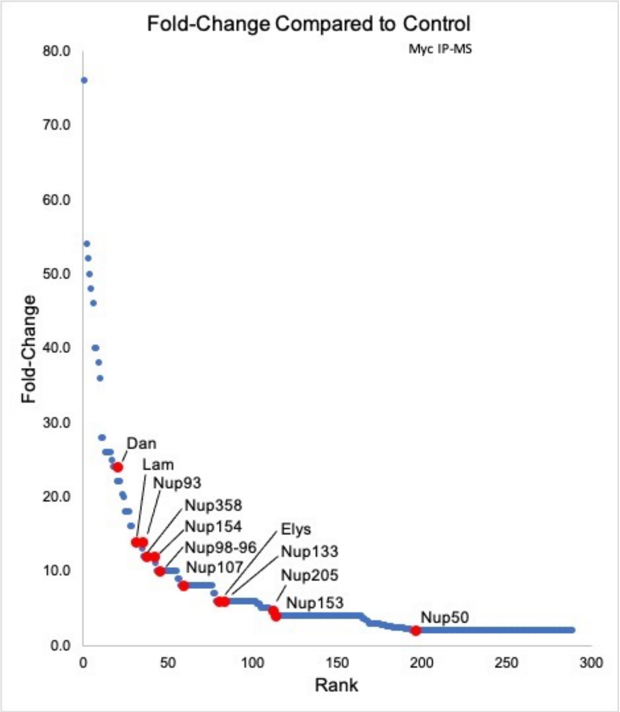
